# Limited involvement of Ca^2+^/calmodulin-dependent kinase II α and β in mammalian sleep

**DOI:** 10.1101/2024.11.17.624045

**Authors:** Weiwen Yang, Jingyi Shi, Chenggang Li, Jingqun Yang, Jianjun Yu, Juan Huang, Yi Rao

## Abstract

While sleep is important, our understanding of its molecular mechanisms is limited. Over the last two decades, protein kinases have been implicated in sleep regulation, with a prominent role for Ca^++^/calmodulin-dependent kinase II (CaMKII) β. Of all the known mouse genetic mutants, the biggest changes in sleep was reported to be observed in *Camk2b* gene knockout mice: sleep was reduced by approximately 120 minutes (mins) over 24 hours (hrs). We have reexamined the sleep phenotype in *Camk2a* and *Camk2b* knockout mice, and while we have observed sleep reduction in *Camk2a* knockout mice, we did not find sleep reduction in *Camk2b* mutants.We did find both *Camk2a* and *Camk2b* participated in homeostatic sleep rebound, though. Because CamKII α and β are widely known to be crucial kinases with interesting properties, it is worthwhile to keep our results in record for a general service to the field.

## INTRODUCTION

Genetics has been very helpful for our understanding of molecular mechanisms of sleep regulation. Genetic mutants of mice have been particularly useful for revealing roles of specific genes involved in regulating sleep. The most famous mouse mutants with sleep phenotypes are perhaps mutants for orexin and its receptor ^1,2^. They are required for maintaining wakefulness during daytime and loss of function (LOF) mutant mice (or dogs or humans) show characteristic narcoplepsy with cataplexy. Another well-known mouse gene is that encoding SIK3, whose gain of function (GOF) mutation was discovered to increase sleep^3-5^. We have found reduced daytime rapid eye movement (REM) sleep in SIK3 LOF mutants^6^, consistent with the sleep increase phenotype of SIK3 GOF mutant mice^3^, although the extent of reduction in LOF mutants was very moderate^6^.

Other protein kinases implicated in sleep include: protein kinase A (PKA)^7-9^, the extracellular signal-regulated kinase (ERK)^10-12^, the adenosine monophosphate (AMP)-activated protein kinase (AMPK)^13-15^, CaMKII α and β^16−18^, c-Jun N-terminal kinase (JNK)^19^, and the liver kinase B (LKB1)^20^. The most prominent changes in sleep was reported in *Camk2b* knockout mice^17^: a reduction of more than 120 minutes (mins) in 24 hours. This change is bigger than changes in all other known mouse mutants^1-6,9,12,18,20-31^.

It thus seemed that *Camk2b* knockout mice are the most important mutant mice for further studies of sleep regulation. We constructed the embryonic knockout mice and tested the sleep behavior after backcross, yet could not validate the result from the functional quick screen reported previously^17,18^.

## RESULTS

### Embryonic knockout of *Camk2a* and *Camk2b*

In our efforts to study protein phosphorylation in sleep, we generated *Camk2a* and *Camk2b* knockout mice (Figure 1A and 1B). While the previous paper reporting sleep reduction used embryonic injection of simple guide (sg) RNAs to create LOF *Camk2a* and *Camk2b* mutants for functional quick screen^17,18^, we used sgRNAs to create genetically transmissible mutants and crossed them for at least six generations before functional testing. We deleted exons 5 and 6 of *Camk2a* which encoded amino acid residues (aa) 92 to 137, with an additional frameshift mutation of CCU (Pro) to CUG (Leu), resulting in truncation thereafter (Figure 1A). The shifted frame covered all aa sequences targetted for deletion in the previous paper^17^. We deleted exons 7 and 8 of *CaMK2b* encoding aa 139 to 200, with an additional frameshift mutation from GGGGUG (Gly-Val) to GGGUGA (Gly-*), which also covered all sequences deleted in the the previous paper^17^. Western blot analysis with antibodies recognizing CaMKIIα and CaMKIIβ proteins confirmed the effectiveness of *Camk2a* and *Camk2b* gene targeting in heterozygous mutant mice and homozygous mutant mice (Figure 1C and 1D). As behavioral controls, we tested *Camk2a* and *Camk2b* knockout mice in Morris water maze and confirmed that they were defective (Supplementary Figure S1 A and B).

**Figure 1.**
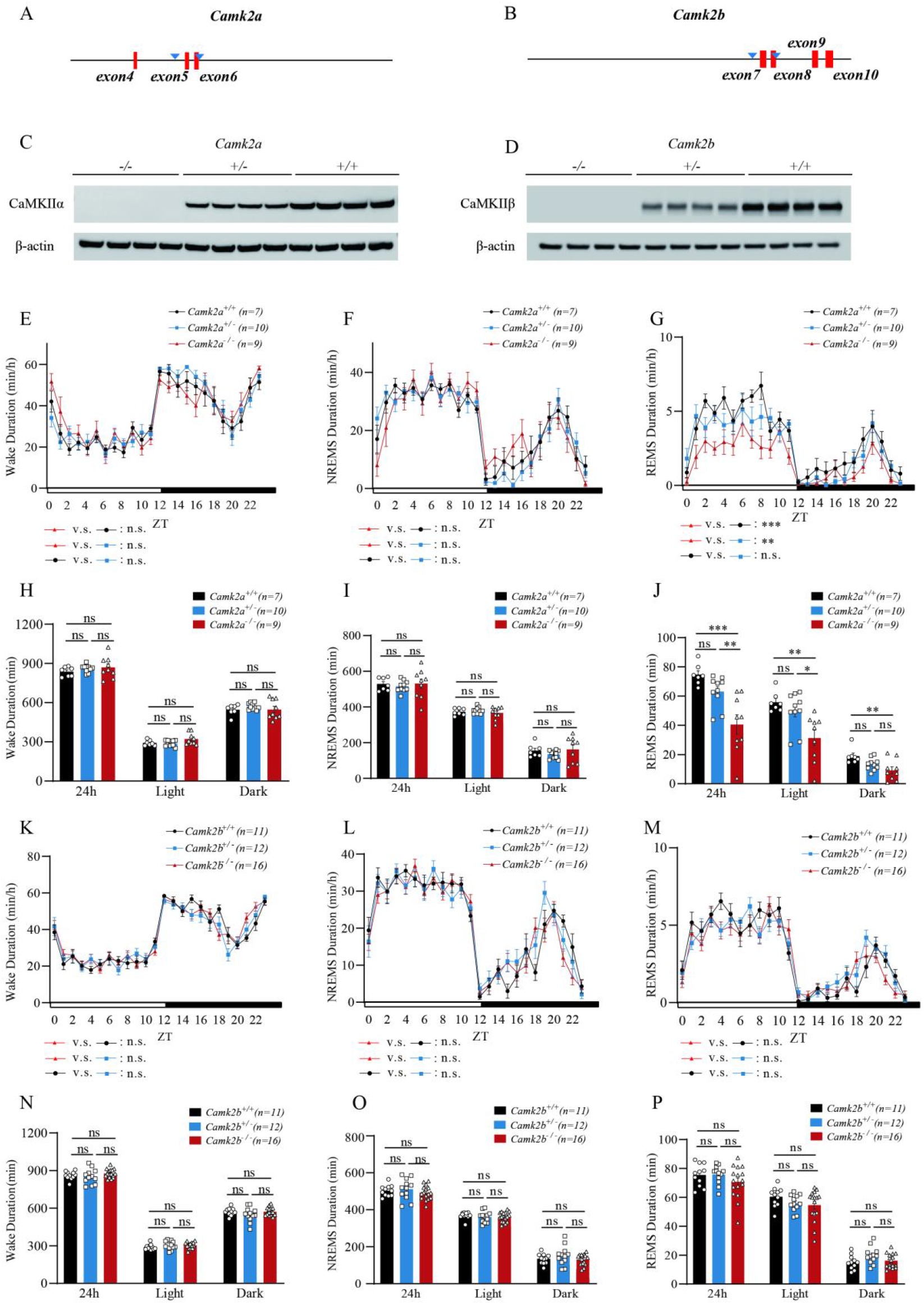
Generation of *Camk2a* and *Camk2b* Mutant Mice and Their Sleep Phenotypes. (**A**) A schematic diagram illustrating the strategy using a pair of sgRNAs to mediate the deletion of exons 5 and 6 in *Camk2a* with sgRNAs CTGGATCACGAAGACCCCTG and ACCTGAAGGTGAGTAACCCT. (**B**) A schematic diagram for the deletion of exons 7 and 8 in *Camk2b* with sgRNAs TTGCAGTCACCTATGTCACG and TGTGTGAGGGGAAACACCTG. (**C**) Western analysis of CaMKIIα from the brain lysates of mice with the genotypes of *Camk2a*^-/-^, *Camk2a*^+/-^ and *Camk2a*^+/+^ (4 mice for each genotype shown here). (**D**) Western analysis of CaMKIIβ from the brain lysates of mice with the genotypes of *Camk2b*^-/-^, *Camk2b*^+/-^ and *Camk2b*^+/+^ (4 mice for each genotype shown here). (**E-J**) Analysis of sleep in *Camk2a*^+/+^ (n = 7, black curve and bar), *Camk2a*^+/-^ (n=10, blue curve and bar) and *Camk2a*^-/-^ (n = 9, red curve and bar) mice. Profiles showing wake time of each hour in min/hr. (**E**), profiles of NREM sleep (**F**), or profiles of REM sleep (**G**). The x axis shows zeitgeber time (ZT) with the white box indicating light phase (or daytime) and black box dark phase (or nighttime). 24 hrs, daytime and nighttime durations of wake (**H**), NREM sleep (**I**) or REM sleep (**J**) shown in mins. **(K-P**) Sleep of *Camk2b*^+/+^ (n=11, black curve and bar), *Camk2b*^+/-^ (n=12, blue curve and bar) and *Camk2b*^-/-^ (n=16, red curve and bar) mice. Profiles of wake time (**K**), NREM sleep (**L**) or REM sleep (**M**). 24 hr, daytime and nighttime durations of wake (**N**), NREM sleep (**O**) and REM sleep (**P**). ns, not significant; *p < 0.05; **p <0.01; ***p <0.001 and ****p <0.0001; mean ± standard error of the mean (mean ± SEM). Two-way ANOVA (E, F, G, K, L, M); One-way ANOVA (H, I, J, N, O, P)

### No significant change was observed for *Camk2b*^-/-^ mice in basal sleep

We then analyzed the sleep phenotypes in mutant mice according to procedures we have published previously^6,20^. In *Camk2a* knockout mice, while no significant change was observed in total wake duration (Figure 1E and 1H), daytime or nighttime non-REM (NREM) sleep (Figure 1F and 1I), REM sleep was found to be reduced in daytime for 25.5 mins and nighttime for 9.7 mins (Figure 1G and 1J, Table 1, Supplementary Figure S2 I and J). These were not very different from the total of approximately 50 mins over 24 hrs reported previously^17^.

**Table 1.** Total Time Spent in Different Sleep-Wake States of *Camk2a* Knockout Male Mice.

| | Total time | 24 hours (Mean ± SEM, min) | Light Phase (Mean ± SEM, min) | Dark Phase (Mean ± SEM, min) |
| --- | --- | --- | --- | --- |
| Sleep | *Camk2a*<sup>+/+</sup> | 603.0±13.9 | 428.6±7.9 | 174.4±15.4 |
| | *Camk2a*<sup>+/-</sup> | 579.1±9.1 | 430.7±7.2 | 148.4±8.1 |
| | *Camk2a*<sup>-/-</sup> | 570.3±27.0 | 397.6±14.7 | 172.6±24.5 |
| NREM | *Camk2a*<sup>+/+</sup> | 528.3±13.7 | 372.8±7.4 | 155.5±8.2 |
| | *Camk2a*<sup>+/-</sup> | 516.0±10.4 | 380.9±5.7 | 135.1±7.3 |
| | *Camk2a*<sup>-/-</sup> | 529.7±26.9 | 366.3±11.5 | 163.4±23.7 |
| REM | *Camk2a*<sup>+/+</sup> | 74.7±2.7 | 55.8±2.6 | 18.9±2.0 |
| | *Camk2a*<sup>+/-</sup> | 63.1±3.3 | 49.8±4.0 | 13.3±1.3 |
| | *Camk2a*<sup>-/-</sup> | 40.5±6.6 | 31.3±5.3 | 9.2±2.4 |
| Wake | *Camk2a*<sup>+/+</sup> | 837.0±13.9 | 291.4±7.9 | 545.6±15.4 |
| | *Camk2a*<sup>+/-</sup> | 860.9±9.1 | 289.3±7.2 | 571.6±8.1 |
| | *Camk2a*<sup>-/-</sup> | 869.7±27.0 | 322.4±14.7 | 547.4±24.5 |

However, when we analyzed the sleep phenotype of *Camk2b* knockout mice (Figure 1K to 1P, Table 2, Supplementary Figure S2C, S2D, S2G, S2H, S2K, S2L), no significant change in sleep was observed: from total wake duration, to total sleep, daytime or nighttime NREM sleep or REM sleep.

**Table 2.**
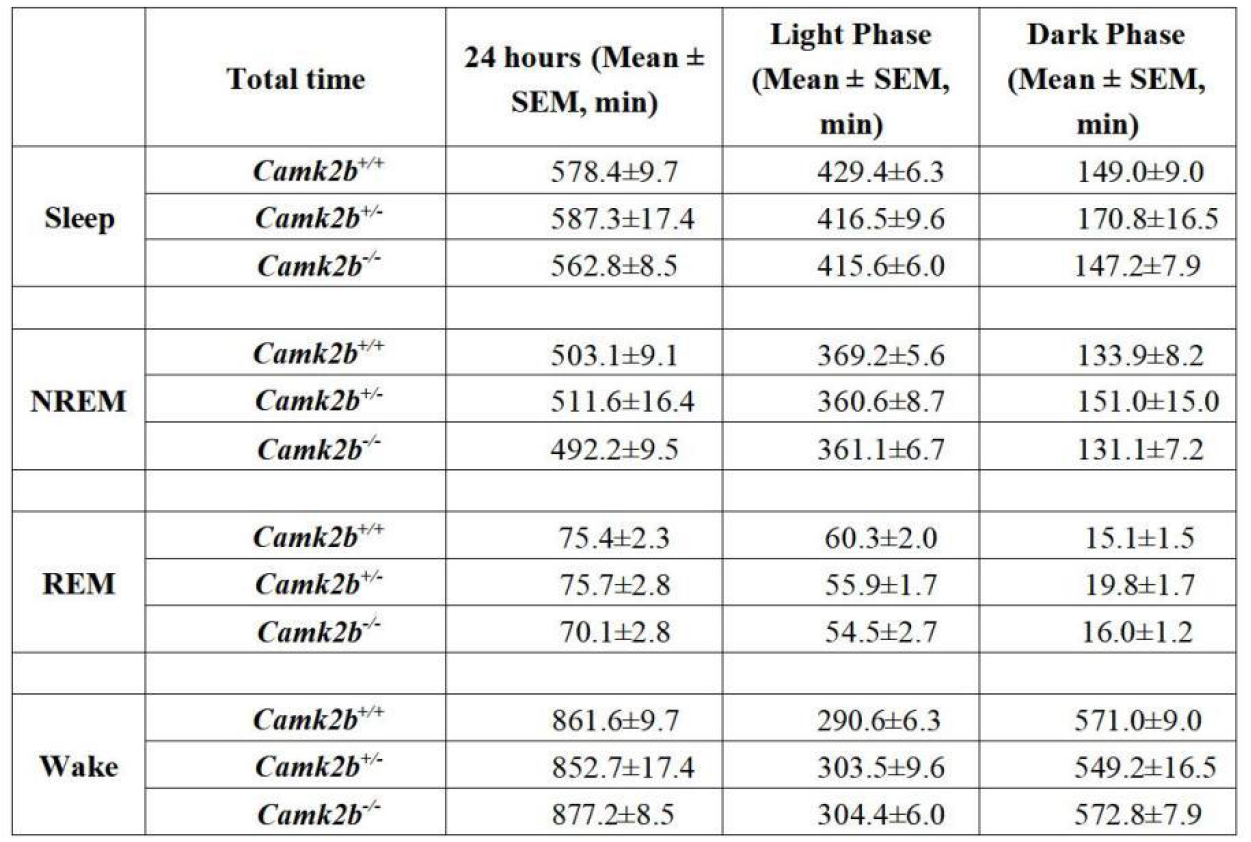
Total Time Spent in Different Sleep-Wake States of *Camk2b* Knockout Male Mice.

### Camk2a and camk2b did participate in homeostatic sleep rebound

We also conducted sleep deprivation (SD) experiment and analyzed the rebound after 6 hrs of SD in both mutant mice. In *Camk2a* knockout mice, there were no significant alterations in the NREM rebound or wake reduction (Figure 2A and 2B). A slight decrease in REM rebound was found after 14 to 20 hours; however, no significant differences were observed 24 hours post-SD (Figure 2C). In *Camk2b* knockout mice, although cumulative NREM rebound and wake reduction were found to decrease slightly after 13 to 17 hours, there were no significant differences in sleep rebound 24 hours after SD comparing to wild-type mice (Figure 2D and 2E). And no differences in cumulative REM rebound were detected (Figure 2F).

**Figure 2.**
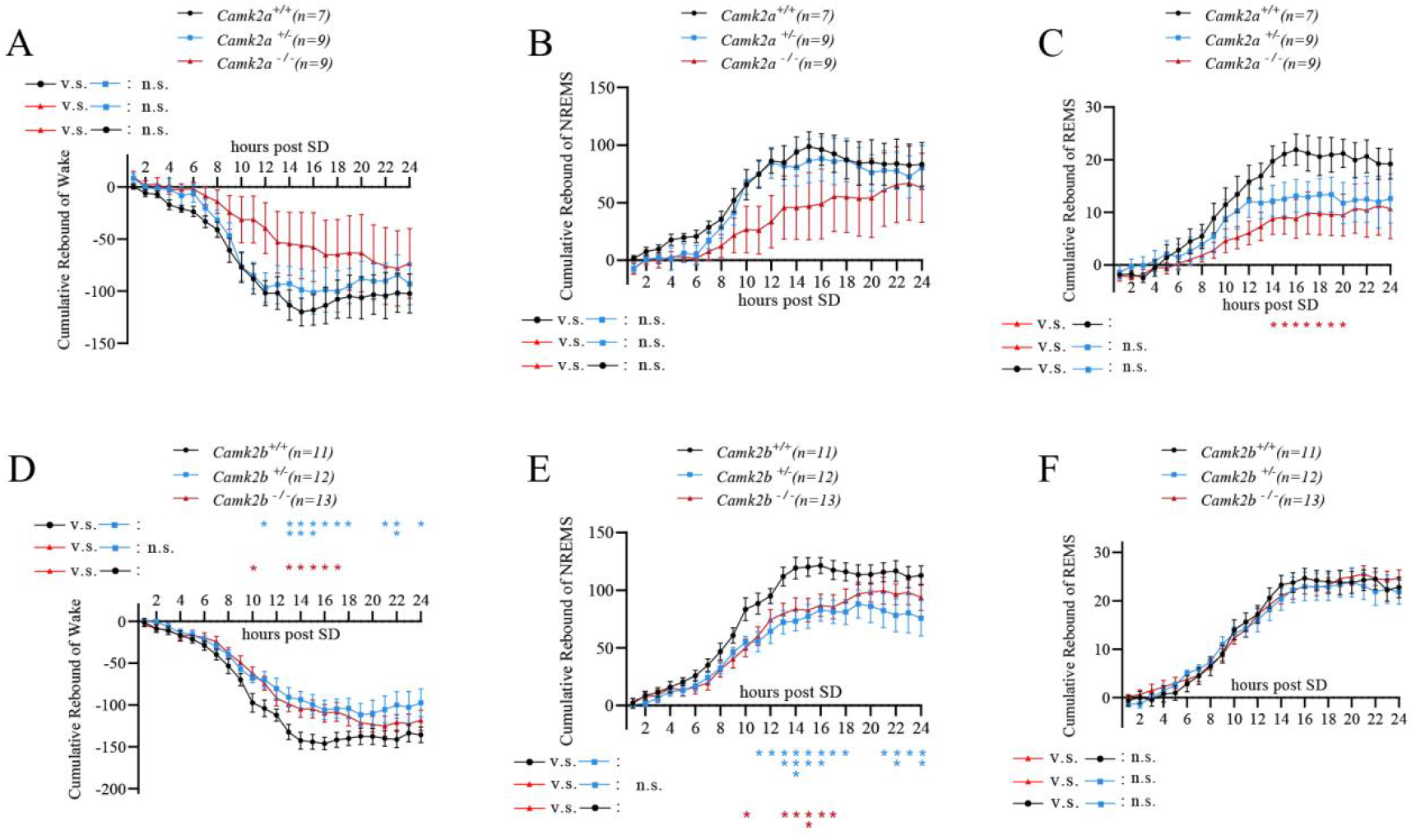
Sleep rebound in *Camk2a*^-/-^ and *Camk2b*^-/-^ mice after sleep deprivation. (A-C) Cumulative wake reduction, and NREM, REM rebound of *Camk2a*^+/+^ (n = 7, black curve), *Camk2a*^+/-^ (n = 9, blue curve) and *Camk2a*^-/-^(n = 9, red curve) after 6 hrs of SD. (D-F) Cumulative wake reduction, and NREM, REM rebound of *Camk2b*^+/+^ (n = 11, black curve), *Camk2b*^+/-^ (n = 12, blue curve) and *Camk2b*^-/-^(n = 13, red curve) after 6 hrs of SD. ns, not significant; *p < 0.05; **p <0.01; ***p <0.001; mean ± standard error of the mean (mean ± SEM). Two-way ANOVA was used (A-F).

## DISCUSSION

Thus, we have confirmed the involvement of CaMKIIα in learning, basal sleep and homeostatic sleep rebound, and that of CaMKIIβ in learning and homeostatic sleep rebound, but did not find a role for CaMKIIβ in basal sleep regulation.

There can be multiple explanations for the discrepancies between the previous results^17^ and our present results. The easier one would be the possibility that, while the previous study had used mutants quickly generated by embryonic injection of sgRNAs and functionally tested in the same generation, we have used the more traditional and reliable method of generating stable mutants which were functionally tested after a few generations. While the quick method has the advantage of time-saving, ours is the reliable method used by the vast majority of users.

## METHODS

### Mouse stocks

WT C57 BL/6J mice (8 to 10 weeks old) were purchased from Beijing Vital River Laboratories Technology Co., Ltd. or Laboratory animal resource center in Chinese Institute for Brain Research. CaMK2α and CaMK2β knock out mice were constructed with CRISPR-Cas9 gene targeting technology. For CaMK2α-KO, the gRNA sequences were designed as 5’-CTGGATCACGAAGACCCCTG-3’ and 5’-ACCTGAAGGTGAGTAACCCT-3’ to delete the 5^th^ and 6^th^ exons. For CaMK2β-KO, the gRNA sequences were designed as 5’-TCTAGGACCTCCATGTTGGG-3’ and 5’-TAAGACCTGTGTGTGAGAGG-3’ to delete the 7^th^ and 8^th^ exons. A mixture of Cas9-expressing mRNA and sgRNAs was injected into fertilized eggs through electroporation and the eggs were then transplanted into the womb of foster mothers. F0 and F1 mice were genotyped through PCR to make sure the presence of recombination. The genotyping primers for CaMK2α-KO were 5’-AGGGGACAGAGAGGGGTAAG-3’ and 5’-ACAGGGCAGCCTCAGTCTAA-3’. For CaMK2β-KO, primers were 5’-CAGGCCTACGATGGAGATGT-3’ and 5’-ATCCTGGTGTCCACTTGCTC-3’. Mutant lines were back-crossed to C57BL/6J for at least 5 generations to exclude possible off-targeting.

### Mouse housing

All experimental procedures were performed in accordance with the guidelines and were approved by the Animal Care and Use Committee of Chinese Institute for Brain Research, Beijing. Mutant mice and wt littermates were maintained on a C57 BL/6J background. Mice were housed under a 12 hr:12 hr light/dark cycle and controlled temperature and humidity conditions. Food and water were delivered *ad libitum*. Mice used in all experiments were 10-14 weeks old.

### EEG and EMG Recording and Analysis

EEG and EMG data recording and analysis were performed as our previous study^20^. EEG and EMG data at basal sleep conditions were recorded for 2 consecutive days, with a sample frequency of 200 Hz and epoch length of 4 seconds. EEG and EMG data were initially processed using AccuSleep^32^ and then were manual correction in SleepSign. EEG and EMG signals were classified into Wake (fast and low amplitude EEG, high amplitude and variable EMG), NREM (high amplitude and 1-4 Hz dominant frequency EEG, low EMG tonus) and REM (low amplitude and 6-9 Hz frequency EEG, complete silent of EMG). The state episode was defined as at least three continuous and unitary state epochs. Epoch contained movement artifacts were included in sleep duration analysis but excluded from the subsequent power spectrum analysis. For power spectrum analysis, EEG was subjected to fast Fourier transform analysis (FFT). Power spectra represents the mean ratio of each 0.25 Hz to total 0–25 Hz of EEG signals during 24 hrs baseline condition. The power density of NREMs represents the ratio of delta power density (1-4 Hz) to total power (0-25 Hz) in each hr. Cumulative rebound represented cumulative changes of time in post-SD compared with relative ZT under the baseline condition. Sleep/wake transition probabilities was analyzed as described in a previous study^33^. For instance, *PW to NR* = N*W to NR* / (N*W to W* + N*W to R* + N*W to NR*), N*W to NR* denotes the number of transitions that transit from wakefulness epoch to NREM sleep epoch. W: wakefulness epoch, NR:NREM epoch, R: REM epoch. The changes of NRMES delta density denotes the difference before and after sleep deprivation, which are calculated as the value of the NRMES delta density after sleep deprivation minus that of the baseline at the equivalent ZT point.

### Morris Water Maze

Spatial learning and memory were assessed using the hidden-platform version of the Morris water maze^34,35^. A circular tank (150 cm in diameter) was filled with opaque water (21 ± 2 °C) to a depth of 35 cm to make the platform invisible. The tank was divided into four quadrants, with the escape platform consistently positioned 1 cm below the water surface in the center of the SE quadrant. Data collection was facilitated by a digital camera connected to an image tracking system. The mice were given 1min of free swimming on the first day. Subsequently, the mice were brought to the water maze room 1h early for acclimatization before each daily session. Over 6 consecutive days, the mice performed four trials per day with an intertrial interval of 30min. The mice started each trial facing the tank wall, and the starting position was randomized. A maximum of 60s was allowed for the mice to locate the platform. After reaching the platform within 60s, the mice were allowed to remain on it for 15s. In cases where the mice failed to locate the platform within 60s, they were gently guided to the platform and allowed to stay on it for 15s. The time taken to reach the platform was analyzed.

### Sleep Deprivation

After 2 consecutive days of EEG and EMG signals recording, mice were introduced into new cages at ZT0 or ZT6. Mice were gently handled or touched to keep them awake for 6 hrs of sleep deprivation, before being returned to the recording cage for another 24 hrs of recording.

### Statistical Analysis

All statistical analyses were performed using GraphPad Prism 9.0. One-way ANOVA was used to compare differences among more than three groups, followed by Tukey’s multiple comparisons tests. Kruskal-Wallis tests were used for non-parameters tests. Two-way ANOVA was used to compare the differences between different groups with different treatments, followed by Tukey’s multiple comparisons tests. Two-way ANOVA with repeated measurements (Two-way RM ANOVA) was used when the same individuals were measured on the same outcome variable more than once, followed by Tukey’s multiple comparisons test. Data are presented as mean±SEM. In all cases, p values more than 0.05 were considered not significant.

## Legends for Supplemental Figures

**Supplemental Figure S1.**
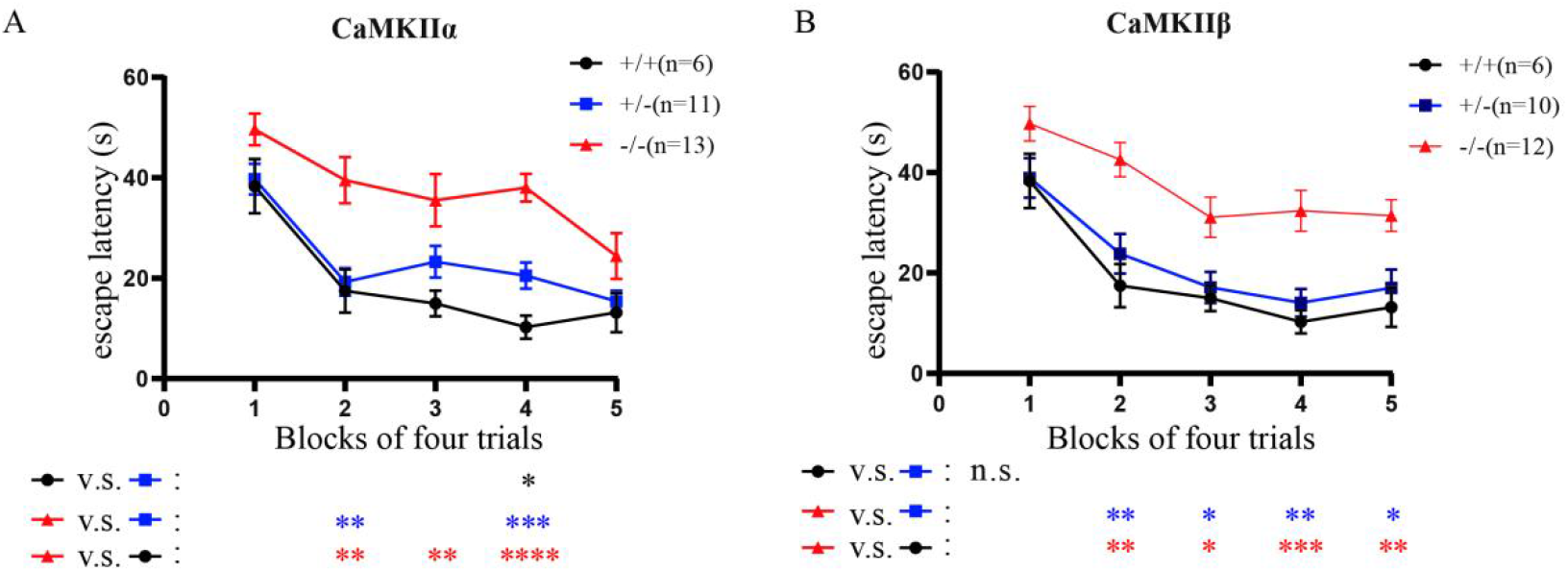
Behaviours of *Camk2a* and *Camk2b* knockout male mice in the Morris water maze. (**A**) Comparison among WT mice (n=6, black curve), *Camk2a*^+/-^ mice (n=11, blue curve), *Camk2a*^-/-^ mice (n=13, red curve). Escape latency in seconds (s) are displayed in the average of four trails daily and the trial lasted 5 days. (**B**) Comparison among *Camk2b*^+/-^ mice (n=10, blue curve) and *Camk2b*^-/-^ mice (n=12, red curve). ns, not significant; *p < 0.05; **p <0.01; ***p <0.001 and ****p <0.0001; mean ± standard error of the mean (mean ± SEM). Two-way ANOVA (A, B).

**Supplemental Figure S2.**
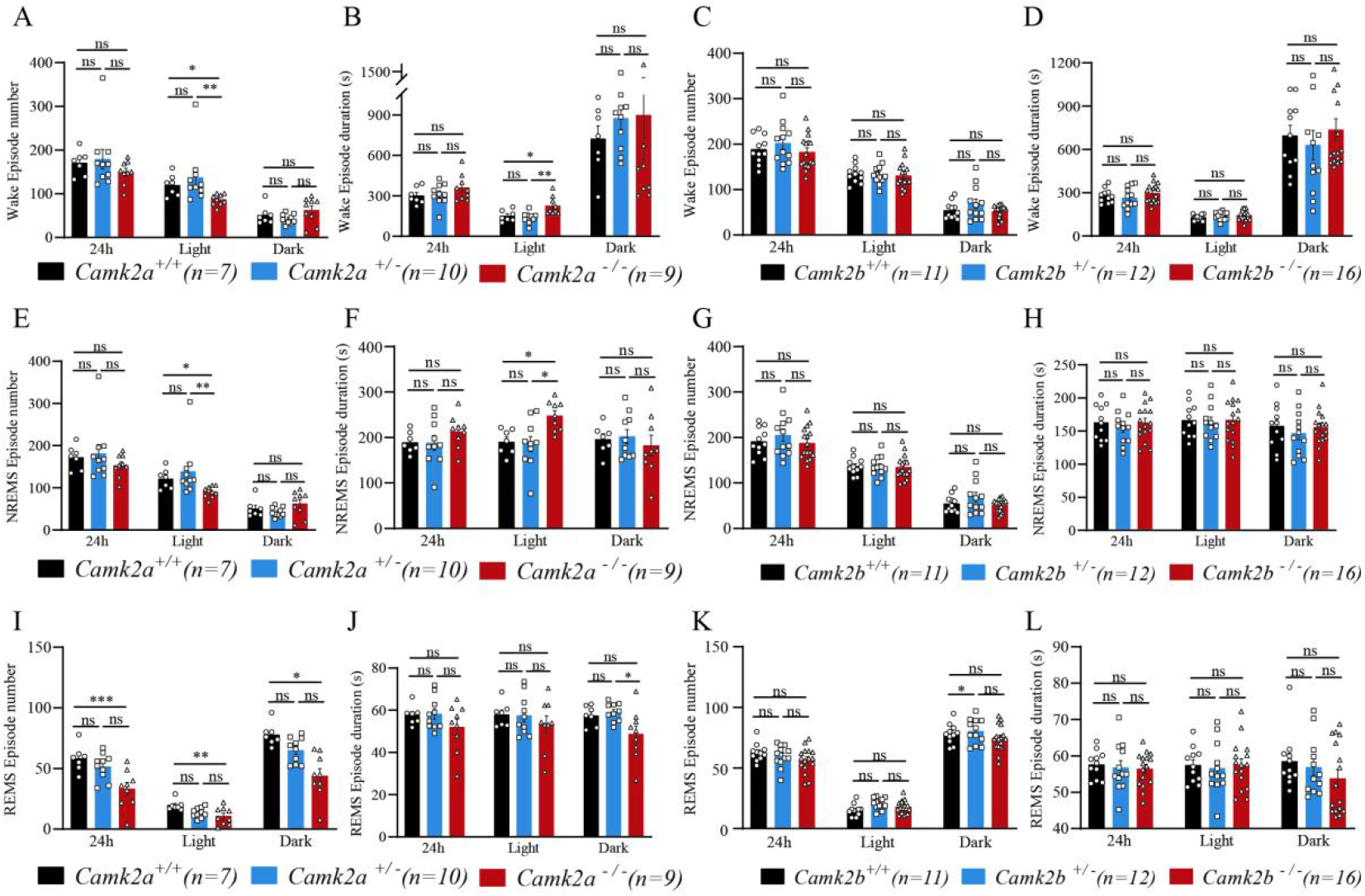
Detailed description of the sleep phenotypes and learning behaviour of *Camk2a* and *Camk2b* knockout male mice. (**A-L**) The wake time, NREM sleep, REM sleep episode number and duration of *Camk2a* and *Camk2b* knockout mice in 24 hr, daytime or nighttime. The wake time episode number (**A**) and duration (**B**) of *Camk2a* knockout mice and the number (**C**) and duration (**D**) of *Camk2b* knockout mice. The NREM sleep episode number **(E**) and duration (**F**) of *Camk2a* knockout mice and the number (**G**) and duration (**H**) of *Camk2b* knockout mice. The REM sleep episode number (**I**) and duration (**J**) of *Camk2a* knockout mice and the number (K) and duration (**L**) of *Camk2b* knockout mice. ns, not significant; *p < 0.05; **p <0.01; ***p <0.001 and ****p <0.0001; mean ± standard error of the mean (mean ± SEM). Kruskal-Wallis test (A-L).

